# Functional stress responses in Glaucophyta: evidence of ethylene and abscisic acid functions in *Cyanophora paradoxa*

**DOI:** 10.1101/2023.06.22.546102

**Authors:** Baptiste Genot, Margaret Grogan, Matthew Yost, Gabriella Iacono, Stephen D. Archer, John A. Burns

## Abstract

Glaucophytes, an enigmatic group of freshwater algae, occupy a pivotal position within the Archaeplastida, providing insights into the early evolutionary history of plastids and their host cells. These algae possess unique plastids, known as cyanelles, that retain certain ancestral features, enabling a better understanding of the plastid transition from cyanobacteria. In this study, we investigated the role of ethylene, a potent hormone used by land plants to coordinate stress responses, in the glaucophyte alga *Cyanophora paradoxa.* We demonstrate that *C. paradoxa* produces gaseous ethylene when supplied with exogenous 1-aminocyclopropane-1-carboxylic acid (ACC), the ethylene precursor in land plants. In addition, we show that cells produce ethylene natively in response to abiotic stress, and that another plant hormone, abscisic acid (ABA), interferes with ethylene synthesis from exogenously supplied ACC, while positively regulating reactive oxygen species (ROS) accumulation. ROS synthesis also occurred following abiotic stress and ACC treatment, possibly acting as a second-messenger in stress responses. A physiological response of *C. paradoxa* to ACC treatment is growth inhibition. Using transcriptomics, we reveal that ACC treatment induces the upregulation of senescence-associated proteases, consistent with the observation of growth inhibition. This is the first report of hormone usage in a glaucophyte alga, extending our understanding of hormone-mediated stress response coordination into the Glaucophyta, with implications for the evolution of signaling modalities across Archaeplastida.

## INTRODUCTION

Blue-green algae belonging to the monophyletic group Glaucophyta constitute an ancient lineage that, along with Rhodophyta and Chloroplastida, is thought to have originated from a primary endosymbiosis between cyanobacteria and a phagotrophic eukaryote (Price et al., 2016). Today, 4 genera (*Cyanophora, Gloeochaete, Cyanoptyche, and Glaucocystis*) and 25 glaucophyte species are known (Guiry and Guiry 2024). They are all unicellular freshwater phototrophs and are notably characterized by non-stacked thylakoids and two unique traits inherited from a prokaryotic ancestor: peptidoglycan layers around the plastid called the muroplast or cyanelle in glaucophytes and carboxysome-like bodies (Jackson et al., 2015; Price et al., 2016). For these reasons, glaucophytes are sometimes called “living fossils,” as they represent one of the earliest diverging lineages within the Archaeplastida and retain traits of the cyanobacterial endosymbiont that are absent in red or green algae.

*Cyanophora paradoxa* is the only glaucophyte species whose nuclear genome has entirely been sequenced (Price et al., 2016, 2012); it has therefore been used as a model glaucophyte in studies of the early evolution of photosynthetic eukaryotes. *C. paradoxa* cells are ovoid (9–16 µm) with two cilia and one to two plastids (Jackson et al., 2015). They can easily be cultivated in basic freshwater media in a light and temperature-controlled chamber, making them a good representative of glaucophyte taxa. These algae are found worldwide in habitats poor or rich in nutrients, such as lakes, swamps, acidic sphagnum bogs, or ditches (Price et al., 2016).

In land plants or embryophytes, largely represented in research by the model plant *Arabidopsis thaliana*, the processing of environmental signals into cellular responses is needed for stress adaptation. These signaling mechanisms are well described, and research is primarily motivated by the need for agricultural innovations by creating new crop varieties able to cope better with a constantly changing environment. Notably, studying how plants use small molecule hormones called “phytohormones” to coordinate developmental processes and stress responses leads to a complex but still incomplete model describing ways to induce plants to sustain growth even under biotic and abiotic stresses.

Hormone signaling is poorly understood in primary plastid-bearing algae, which share a common ancestor with land plants. Hormone research in algae mainly focuses on green algae, with *Chlamydomonas reinhardtii* used as a model species (Yoshida et al., 2003), and a few other green algal species with commercial potential (Hu et al., 2021; Mahadi et al., 2022). To extend our knowledge of hormone signaling in Archaeplastida, we tested hormone production and responses in the glaucophyte *Cyanophora paradoxa.* Iconic phytohormones with a significant role in coordinating land plant developmental and environmental signaling and responses include salicylic acid, jasmonic acid, abscisic acid, ethylene, auxins, cytokinins, gibberellic acid, and brassinosteroids (Asami and Nakagawa, 2018).

From algal genomes, including that of *C. paradoxa*, it is not yet possible to predict whether a given hormone might play an active role in the cell. This is in part because the biosynthesis genes, receptors, and downstream effectors of phytohormones often have poor homology between well-described land plant versions and genes in distantly related algae (Bidon et al., 2020). Even when homology can be inferred, the binding of specific molecules to the algal proteins and the resultant downstream protein-protein interactions cannot yet be computationally predicted accurately in novel contexts (Evans et al., 2022; Mou et al., 2021). One well-known hormone in plants is ethylene (C2H4), the only gaseous molecule described as an active phytohormone (Schaller and Kieber, 2002). In plants, ethylene plays various but essential roles in both stress responses and development (Binder, 2020). It has been intensely studied in the model plant *Arabidopsis* for the last 20 years since the discovery of the first ethylene-insensitive mutant (Thomma et al., 1999). In crops, it can control fruit ripening and is used in postharvest storage control (Cocetta and Natalini, 2022).

Ethylene synthesis in plants is well described and requires only a few steps from the initial amino acid methionine. The first step is the production of S-adenosyl-L-methionine (SAM) from methionine by the enzyme SAM synthetase (SAMS). SAM is a vitally important molecule for methylation processes in cells but is also an essential ethylene precursor. In ethylene signaling, SAM is converted into 1-aminocyclopropane-1-carboxylic acid (ACC) by ACC-synthase (ACS). The final step involves the enzyme ACC-oxidase (ACO), which converts ACC to gaseous C2H4. ACS activity is considered the rate-limiting step of ethylene biosynthesis (Van de Poel and Van Der Straeten, 2014). Therefore, the ACS product and direct ethylene precursor ACC, which is water soluble, is frequently applied in controlled experiments to trigger ethylene synthesis and investigate its downstream effects (Adams and Yang, 1979; Li and Guo, 2018; Park et al., 2023).

Although it is often regarded as a phytohormone, primarily understood for its roles in plant development and stress responses, ethylene production has also been observed outside Embryophytes (Tsavkelova et al., 2006). At least four biological mechanisms are known for ethylene production: the recently discovered mechanism via (2-methylthio)-ethanol in soil bacteria (North et al., 2020), via 2-oxoglutarate in bacteria and fungi (Herr and Hausinger, 2018), via 2-keto-4-methylthiobutyric acid (KMBA) in bacteria and fungi (Nazli et al., 2003), and via the above mentioned 1-aminocyclopropane-1-carboxylate (ACC) in plants and reportedly amoebozoans (Amagai and Maeda, 1992; Pattyn et al., 2021).

Among photosynthetic algae, comparative genomic analyses found homologs of the ethylene biosynthesis genes *ACS* and *ACO*, as well as homologs of the plant signal transduction genes ETHYLENE RESPONSE 1 (*ETR*) and CONSTITUTIVE TRIPLE RESPONSE 1 (*CTR1*) in Zygnematophyceae, the algal lineage sister to land plants, with additional evidence of homologs in unicellular chlorophyte algae like *Chlamydomonas reinhardtii* (Feng et al., 2023). Additionally, zygnematophyte algae can produce ethylene when provided with exogenous ACC (Ju et al., 2015). Moreover, ethylene production has been observed in the chlorophyte green algae *Chlamydomonas reinhardtii* and *Haematococcus pluvialis* (Maillard et al., 1993; Yordanova et al., 2010) and the red algae *Pterocladiella capillacea* and *Gelidium arbuscula* (Garcia-Jimenez et al., 2013). While these studies point to a common ancient origin of ethylene synthesis and responses in Archaeplastida algae, a demonstrable recent divergence of plant ethylene synthesis genes (D. Li et al., 2022) and innovations in ethylene signal transduction (Mao et al., 2022) make it difficult to identify the precise origins and evolution of ethylene signaling.

For this study, we tested the ability of the glaucophyte alga *Cyanophora paradoxa* to convert exogenous ACC to ethylene and measured downstream responses to ACC and abiotic stressors. We observed conversion of the plant ethylene precursor ACC to ethylene, native production of ethylene after stress, and shared transcriptional responses between glaucophytes and plants, consistent with prior observations (Ferrari and Mutwil, 2020), but extending them to specific molecular mediators like ACC and ethylene. Our experimental investigation of ethylene signaling components allows us to extend our functional knowledge of ethylene signaling into a new lineage, the Glaucocystophycea.

## MATERIALS AND METHODS

### Algae strain and culture

*C. paradoxa* CCMP329 was used for this study. Cells were cultivated in a modified AF6 media at 16°C in a Percival culture chamber with 12 hours of daylight (40 µE/m^2^/s). Modified AF6 media was made without ammonia but with 3.85 mM NaNO3 and trace metals following Kropat et al. (2011).

### ACC treatment

ACC (1-aminocyclopropane-1-carboxylic Acid - Sigma 149101) was solubilized in water (200 mM stock) and used to treat *C. paradoxa* cells in the following experiments at the mentioned concentrations.

### Gas Chromatography-Flame Ionization Detector (GC-FID)

For this experiment, 15 mL of *C. paradoxa* cells in culture media were sealed for 48 hours in 15 mL headspace vials with appropriate treatments. Ethylene concentrations in the headspace air (2 mL) were determined by gas chromatography. The SRI 8610C gas chromatograph was equipped with a flame ionization detector (GC-FID) and a 3’ x ^1^/8” HAYESEP D packed column. Headspace gas was injected into the column through a 1000 µL sample loop, with N2 as the carrier gas at a flow rate of 15 mL/min, and the flame gases H2 and air at 25 and 250 mL/min, respectively. The isothermal column oven temperature program was set at 50 °C. Under these conditions, ethylene retention time was ∼2.466 minutes. A commercial ethylene standard (GASCO, USA, 2 ppm C2H4 in N2), diluted sequentially in air, was used to generate the calibration of ethylene concentration and peak area. Cell concentration was determined using a hemocytometer, and results were expressed as ng ethylene/10^6^ cells measured from the 1 mL headspace of the vials. All experiments were repeated three times using the same conditions.

### Experimental treatments: Stress response and ABA treatment

Flg22 peptide (RP19986, GenScript) was solubilized in water and used at a final concentration of 100 nM. CuCl2 (Sigma 307483) was solubilized in water and used at a final concentration of 10 µM. For osmotic shock, cells were pelleted at 500 g for 5 minutes and resuspended in 5 mL AF6 media supplemented with 0.1 M sucrose. After 15 minutes of incubation at 16°C, cells were pelleted again and resuspended in 15 mL fresh AF6 media. For UV-B exposure, cells were incubated 2 hours in a dark box containing a UVB100 (ExoTerra) lamp providing about 115 µW/cm^2^ UV-B radiations (from the manufacturer’s manual). ABA (Sigma 862169) was solubilized in DMSO and used at a final concentration of 100 µM.

### Hydrogen peroxide quantification and visualization

Hydrogen peroxide in *C. paradoxa* cells was quantified using the ROS-Glo H2O2 assay (Promega). 20 mL cultures were treated with different ACC concentrations or a mock (water) and incubated for three days in a culture chamber. Cells were then pelleted for 5 minutes at 500 g and resuspended in 200 µL AF6 media. For each condition, 40 µL resuspended cells quadruplicates were used. The ROS-Glo assay was then conducted following the manufacturer’s instructions, and results were acquired on a Filter-max-F5 plate-reader (Molecular Devices). For ROS visualization in live cells, CellROX Green Reagent (Invitrogen) was used following the manufacturer’s instructions. Cells were observed using a Leica DMi8 inverted epifluorescence microscope. ROS were visualized using excitation at 480 nm and a 510 nm emission filter when using CellRox. Images were captured with a DFC9000 sCMOS camera. All experiments were repeated three times using the same conditions.

### RNA extraction

RNA extraction for RNAseq was achieved using a combination of TRIzol (Invitrogen) and PureLink RNA Mini kit (Invitrogen). *C. paradoxa* cells were pelleted at 600 g for 10 minutes. Subsequently, the cells were lysed in 1 mL TRIzol, and the aqueous phase was separated by adding 0.2 mL chloroform, followed by centrifugation at 12000 g for 5 minutes. The resulting aqueous phase, containing the cell lysate, was loaded onto a PureLink RNA Mini kit column. The RNA extraction was then conducted according to the instructions provided by the manufacturer, yielding purified RNA samples for further analysis.

### RNAseq analyses

Total RNA was submitted to a sequencing facility, Genewiz (Azenta, South Plainfield, NJ), for short-read sequencing on the Illumina HiSeq platform. Poly-A selection was completed prior to library prep at the facility, and cDNA library sequencing was completed with 2 x 150 bp paired-end reads per fragment/template. Sequencing generated an average of 23,645,698 read pairs per sample with triplicate samples per condition. Reads were aligned to the *C. paradoxa* reference transcriptome derived from the reference genome (Price et al., 2019) (Transcriptome file: *Cyanophora paradoxa* CCMP329: Cyapar1_GeneCatalog_transcripts_20200807.nt.fasta.gz) using the program Salmon (v1.1.0) (Patro et al., 2017) with decoys generated from the genome assembly. Quantitative differential expression analyses were performed in R (R Core Team, 2013). Salmon quantification files were imported using the function “tximport” (Soneson et al., 2015). Differential expression was completed with edgeR (v3.40.2) using a generalized linear model, a fold-change cutoff of 1.5, and an FDR of 0.05, such that a DE gene had to be 1.5 times over- or under-expressed compared to the control with an FDR < 0.05 in order to register as differentially expressed. For functional analyses, proteins were re-annotated using the most recent version of InterPro (interproscan v5.59-91.0) (Paysan-Lafosse et al., 2023) and gene ontology (GO) terms assigned by interproscan were used for functional enrichment analyses using the program topGO (v2.50.0; GO.db v3.16.0) (Alexa and Rahnenfuhrer, 2023). Enrichment analyses were visualized with GOplot (Walter et al., 2015).

## RESULTS

### *C. paradoxa* converts ACC into gaseous ethylene

The conversion of ACC to ethylene is a hallmark of plant ethylene synthesis. Exposing plants to exogenous ACC induces the production of ethylene gas (Zhang and Wen, 2010) and the activation of ethylene signaling (Park et al., 2023). To probe whether glaucophyte algae also convert ACC to ethylene, we added ACC to *C. paradoxa* cultures and measured ethylene production by GC-FID. *C. paradoxa* cells treated with ACC produced ethylene, while mock-treated samples did not (Figure 1A). Levels of ethylene gas corresponded to the concentration of added ACC; 0.5, 1, and 5 mM ACC treatments corresponded, respectively, to 49, 137, and 283 ng ethylene/10^6^ cells measured in the headspace of the vials, on average. These data indicate that *C. paradoxa* cells can convert ACC into gaseous ethylene, suggestive of a plant-like ethylene signaling pathway in glaucophytes.

**Figure 1:**
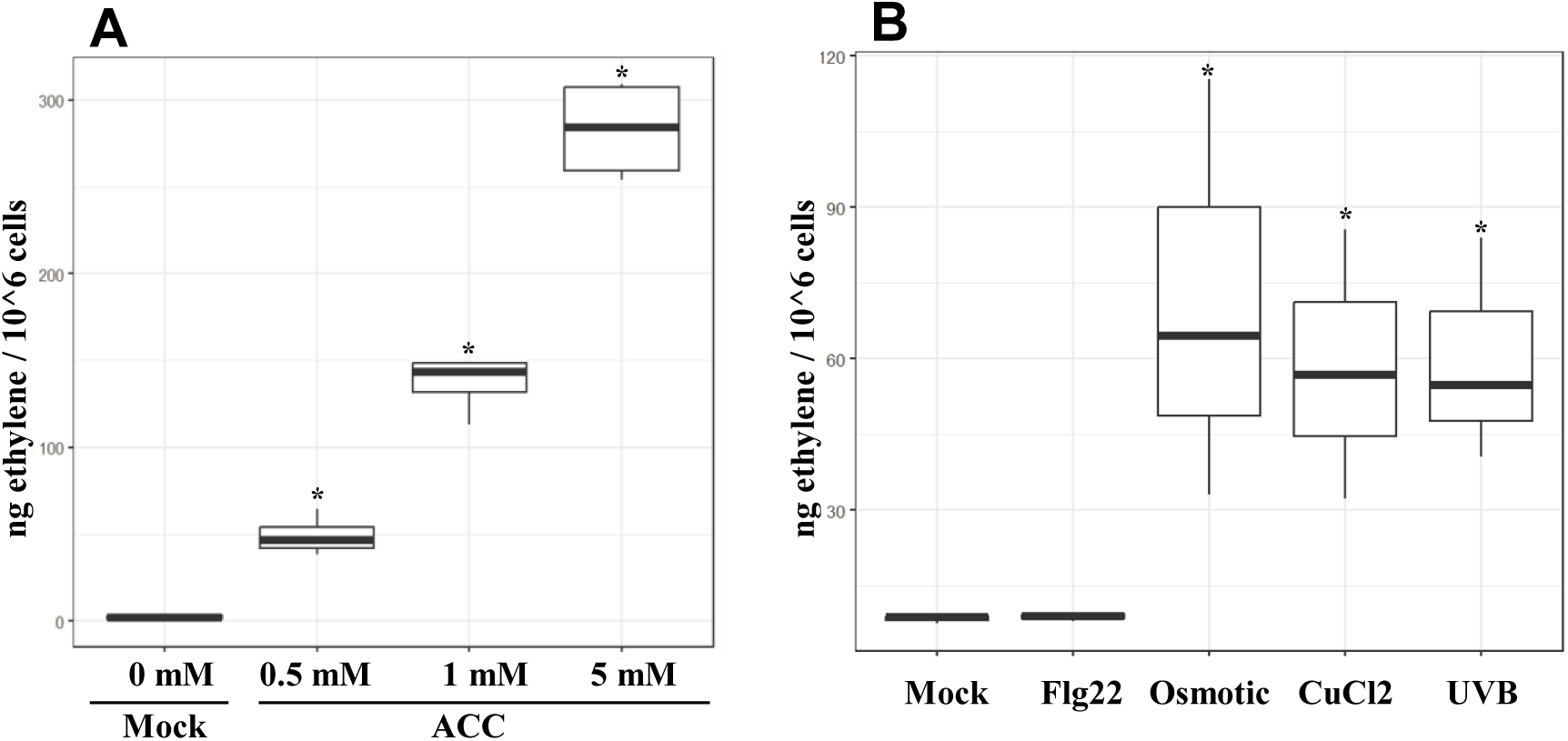
*C. paradoxa* cultures convert ACC to ethylene and produce ethylene natively after stress. Ethylene measurements from *C. paradoxa* cultures using gas chromatography (GC-FID). Results are expressed in ng/ethylene/million cells as measured from the air in the headspace of the vials. To obtain ethylene quantities, the peak area was measured, and ethylene concentrations were determined using a commercial ethylene (Gasco) standard. (A) Samples were treated with ACC before being incubated in sealed vials (*n* = 4 for each treatment) for gas chromatography. All ACC-treated samples displayed significant ethylene production (\**p* < 0.05, ANOVA) compared to mock (water) samples. (B) Samples were either treated with 100 nM flg22 or 10 µM CuCl2, exposed to UV-B for 2 hours, or shocked in 0.1 mM sucrose for 15 minutes before being incubated in sealed vials for gas chromatography (*n* = 3 for each treatment). Asterisks (*) indicate significant ethylene production (\**p* < 0.05, ANOVA) compared to mock (water) treatment.

### *C. paradoxa* makes ethylene natively in response to abiotic stress

As a plant hormone, ethylene is known to regulate plant growth and specific developmental processes (Schaller and Kieber, 2002). It also acts in response to various biotic and abiotic stresses, such as pathogen invasion (Broekaert et al., 2006), chewing insects (von Dahl and Baldwin, 2007), UV-B stress (He et al., 2011), heavy metal stress (Keunen et al., 2016), and osmotic stress (Tao et al., 2015). To test whether *C. paradoxa* cells produce ethylene in response to stresses, *C. paradoxa* cultures were exposed to the synthetic peptide flg22, which mimics bacterial invasion in plants (Meindl et al., 2000), UV-B, CuCl2 oxidative stress, and sucrose-induced osmotic shock. All evaluated abiotic stress conditions (Figure 1B) elicited ethylene production by *C. paradoxa* cultures (∼ 60 ng/10^6^ cells measured in the headspace of the vials, on average) except the candidate biotic stressor, flg22. These results indicate that ethylene synthesis by *C. paradoxa* cells is induced in response to abiotic stress, in agreement with plant stress responses, but not by the biotic stressor flg22, in contrast to plants.

### ROS accumulation occurs following abiotic stress and ACC treatment in *C. paradoxa*

Reactive Oxygen Species (ROS) are common second messengers in signaling pathways, accumulating in response to stress or perception of signaling molecules and contributing to downstream effects within a cell by amplifying the primary signal and activating additional effectors (Schieber and Chandel, 2014). In plants, abiotic stress and ethylene pathway activation are known inducers of ROS accumulation, which is followed by downstream responses like the hypersensitive response, a rapid cell death mechanism aiming to prevent pathogens infections, and/or programmed cell death.

We tested whether abiotic stresses or ACC exposure, both of which induce ethylene production, was followed by ROS accumulation in *C. paradoxa* cells by quantifying hydrogen peroxide (H2O2) content following the treatment (Figure 2A). Osmotic shock and UV-B stress caused a significant increase (*p* < 0.05, Student’s *t*-test) of 8 and 1.6 fold, respectively, in the H2O2 content of *C. paradoxa* cells. Copper-treated cells did not significantly increase H2O2 content (*p* = 0.06, Student’s *t*-test) in these experiments.

**Figure 2:**
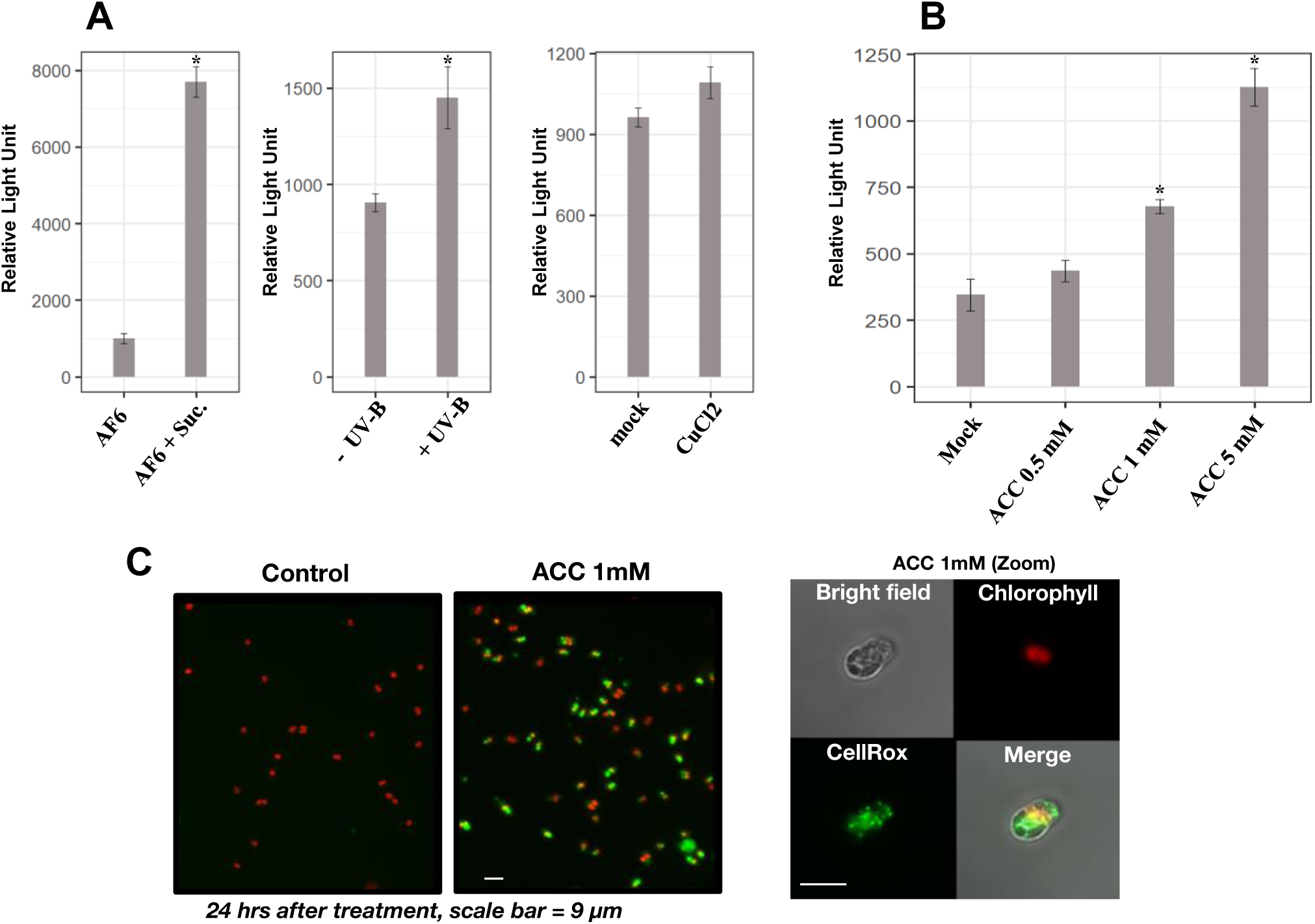
Hydrogen peroxide accumulation is induced downstream of stress and ACC treatments by *C. paradoxa*. (A) Hydrogen peroxide assay (ROS-GLO). Cells were treated the same way as in Figure 1B but were incubated in a culture chamber for 16 hours before being collected for the ROS-GLO assay (*n* = 4 for each condition). Asterisks (*) indicate significantly higher hydrogen peroxide compared to control cells (* *p* < 0.05, Student’s *t*-test). (B) Cells were treated with different ACC concentrations and were incubated in a culture chamber for 24 hours before being collected for the ROS-GLO assay (*n* = 4 for each condition). Asterisks (*) indicate significantly higher hydrogen peroxide compared to control cells (\**p* < 0.05, ANOVA). (C) Fluorescence micrographs of *C. paradoxa* cells treated with 1 mM ACC and CellRox dye allowing ROS visualization. Pictures were taken 24 hours after treatments. Scale bar = 9 µm. Left panel: Control cells; Right panel: ACC-treated cells. Fluorescence channels: Red (Chlorophyll) Ex/Em 650/700 nm; Green (CellRox) Ex/Em 485/520 nm. ACC 1mM (Zoom) picture panel: Detailed view of a *C. paradoxa* cell treated with 1 mM ACC. Upper left: bright field; Upper right: chlorophyll autofluorescence; Bottom left: Cell Rox (ROS); Bottom right: merge of all channels.

Treatment of *C. paradoxa* cells with ACC (Figure 2B) induced a significant accumulation of H2O2 (*p* < 0.05, ANOVA) at treatment concentrations of 1 mM and 5 mM, and the H2O2 content of the cultures positively corresponded to the ACC dosage. We also visualized ROS accumulation (Figure 2C) in *C. paradoxa* cells following ACC treatment using CellRox, a ROS-specific dye. The addition of 1 mM ACC resulted in multiple ROS foci spread through the cells 24 hours after treatment. To test whether ROS accumulation after ACC treatment can be linked more directly to ethylene production, we also tested the effect of ethephon (2-chloroethylphosphonic acid), a commercially available plant growth regulator. When dissolved in an aqueous medium, ethephon spontaneously converts to ethylene, acting as a synthetic ethylene substitute (Lee et al., 2021; Zhang and Wen, 2010). We find that ethephon exposure also induces a strong ROS response (Suppl. Figure 1), suggesting detection of ethylene is a direct mediator of downstream responses.

### ACC treatment causes growth inhibition of *C. paradoxa*

In plants, one major physiological response to the perception of ethylene is the suppression of growth in root and shoot epidermis cells (Vaseva et al., 2018). One reason for growth suppression may be the conservation of resources during stress (Huot et al., 2014). If *C. paradoxa* perceives ethylene as a stress signal, it could similarly enter a resource conservation mode and slow or stop growth. Therefore, we tested the effect of ACC treatment on the growth of *C. paradoxa* (Figure 3). The growth of *C. paradoxa* cultures was monitored during continuous exposure to ACC. Cultures exposed to 1 mM or more ACC exhibited significant growth inhibition by day 5 (*p* < 0.05, ANOVA) (Figure 3). Importantly, when ACC was removed from growth-inhibited cultures, growth resumed, indicating that the growth inhibition is reversible (Suppl. Figure 2). Ethephon treatment also caused growth inhibition (Suppl. Figure 1), suggesting that ethylene perception could be the direct cause of the observed effect.

**Figure 3:**
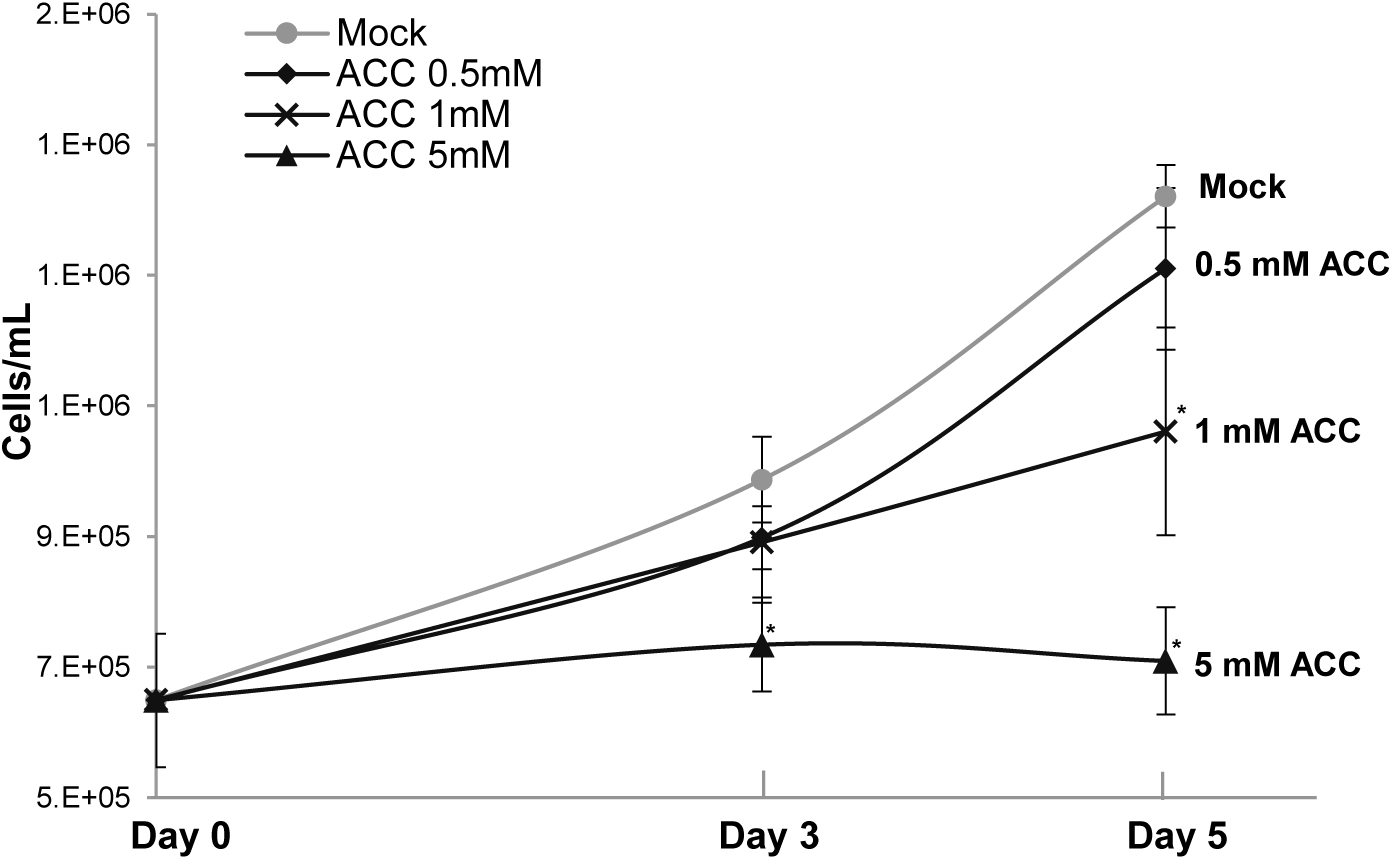
ACC treatment causes growth inhibition of *C. paradoxa* cultures. *C. paradoxa* cell concentrations over 5 days after ACC treatments. 20 mL cultures of *C. paradoxa* were treated with ACC or mock (water) and incubated in a culture chamber for 5 days. Cell concentrations were monitored on days 0, 3, and 5 using a hemocytometer. Asterisks (*) indicate a significant difference compared to control cells at the appropriate time point (\**p* < 0.05, ANOVA).

### ABA interacts with ACC/ethylene signaling in *C. paradoxa*

In light of findings showing that perception and signaling systems for abscisic acid (ABA), another significant phytohormone, are present within algae (Sun et al., 2020), we investigated the possibility of interaction or communication between ABA and ethylene signaling in *C. paradoxa*. In plants, ABA often modulates or inhibits the signaling activities associated with ethylene (Dong et al., 2016).

Cells treated simultaneously with ABA and ACC and assayed after 24 hours exhibited reduced ethylene production by about 50% compared to cells treated with ACC only, indicating that ABA negatively regulates the conversion of ACC to ethylene in *C. paradoxa* (Figure 4A). Individual treatments with either ACC or ABA induced H2O2 production to similar levels (*p* > 0.05, ANOVA) (Figure 4B). Dual treatment with ABA and ACC exhibited an additive effect, with significantly increased H2O2 accumulation compared to the single treatments (*p* < 0.05, ANOVA).

**Figure 4:**
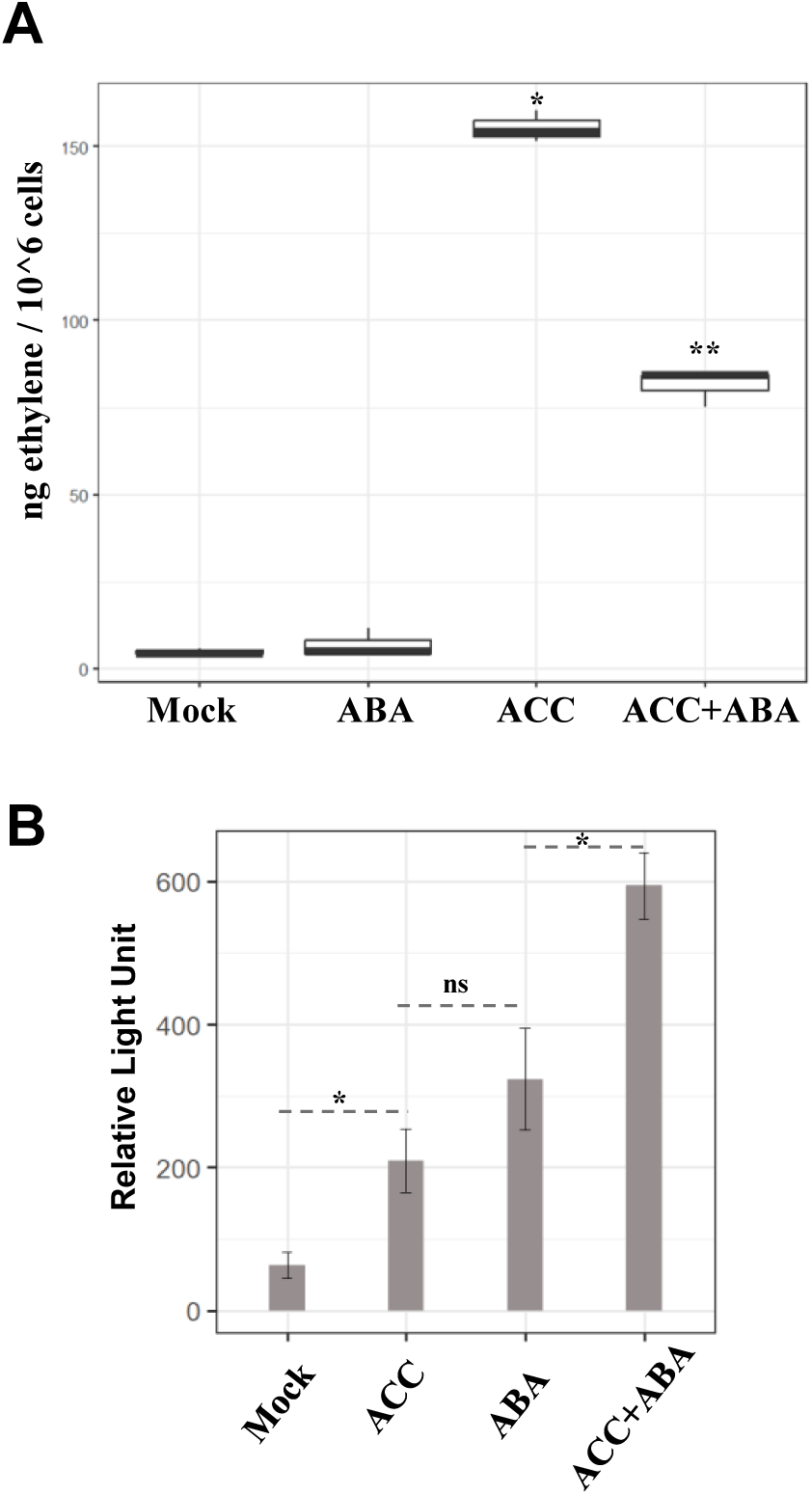
Crosstalk between ACC and ABA signals in *C. paradoxa*. (A) Ethylene measurements using a gas chromatography setup (GC-FID). Results are expressed in ng/ethylene/million cells. Samples were either treated with mock (DMSO), 100 µM ABA, 1 mM ACC, or a combination of ACC and ABA before being incubated in sealed vials for gas chromatography (*n* = 3 for each condition). (*) indicates a significant difference (\**p* < 0.05, ANOVA) compared to the mock treatment. (**) indicates a significant difference between mock and ACC conditions. (B) Hydrogen peroxide assay (ROS-GLO). Cells were treated with different ACC or ABA concentrations and incubated in a culture chamber for 24 hours before being collected for the ROS-GLO assay (*n* = 3 for each condition). (*) indicates a significant difference (\**p* < 0.05, ANOVA) between the two conditions. ns: non-significant differences (*p* > 0.05, ANOVA).

### ACC treatment induces upregulation of senescence-associated proteases in *C. paradoxa*

One output of signal transduction is the alteration of gene expression levels in response to a specific signal or stress. To reveal the glaucophyte response to ACC/ethylene and compare it to that from specific stressors, we conducted RNAseq experiments (Figure 5, Suppl. File 1) to identify differentially expressed genes (DEG) in response to ACC and abiotic stresses (UV-B, osmotic stress, and Cu-induced oxidative stress). After 24 hours of treatment, ACC treatment induced a small number of transcriptional changes, 37 DEGs (28 up and 9 down), despite having large effects on growth (e.g., Figure 3). A low level of copper exposure induced a similarly low number of DEGs (10 up and 4 down). In contrast, cells exposed to pronounced abiotic stress exhibited high numbers of DEGs: (774 up and 1568 down) 24 hours after a 2-hour UV-B stress, and 833 DEGs (388 up and 445 down) after a 15-minute osmotic stress.

**Figure 5:**
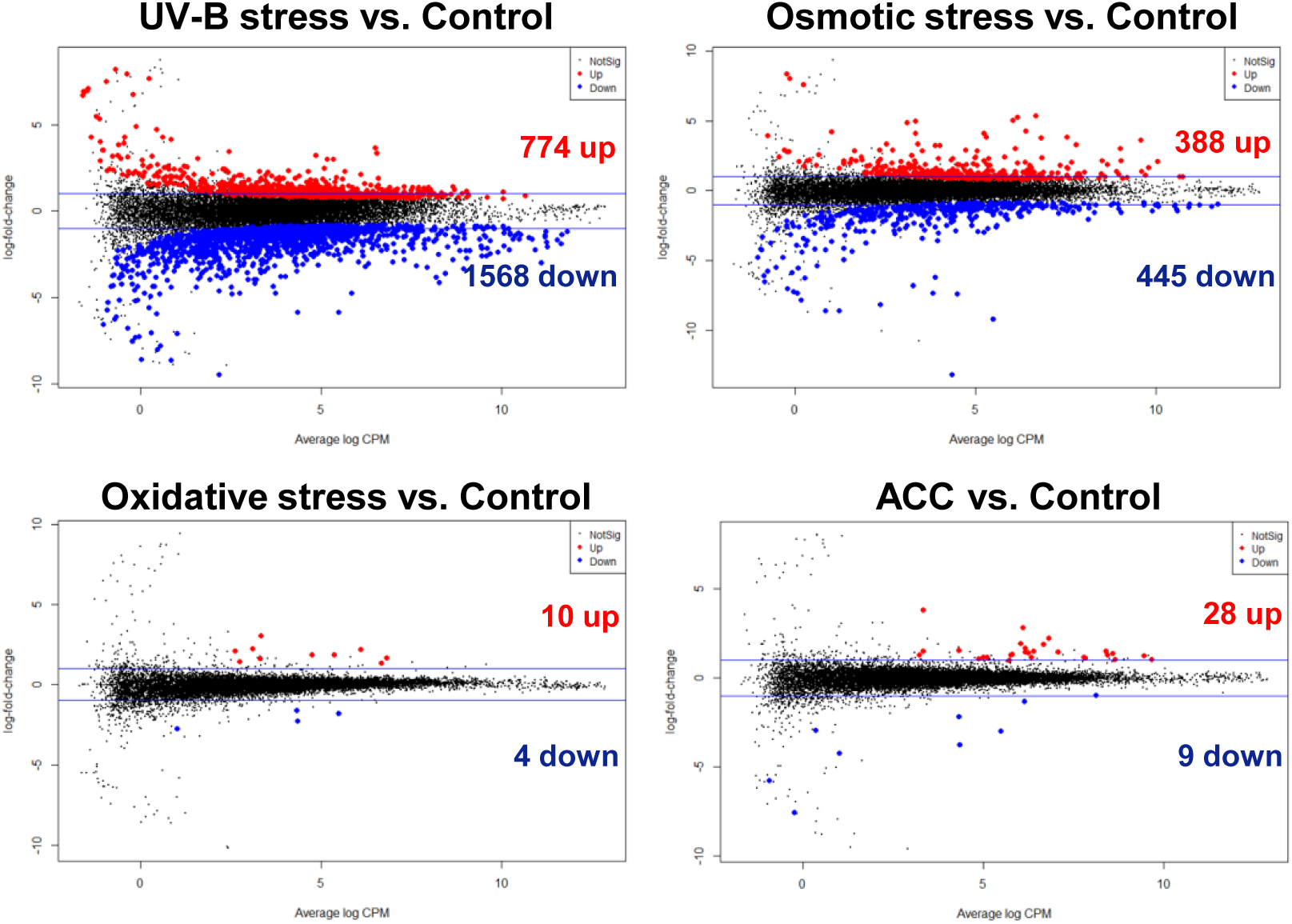
Transcriptional changes after abiotic stress and ACC treatments in *C. paradoxa*. Differentially expressed genes in *C. paradoxa* after abiotic stresses (UV-B, osmotic, and oxidative) and 1 mM ACC treatments. Mean-difference plots (Log fold change vs. Average log CPM) show genes significantly differentially expressed (FDR < 0.05) with a log fold change < log2 (1.5). Upregulated genes are highlighted in red, and downregulated genes are highlighted in blue.

Functional enrichment analysis of DEGs (Figure 6, Suppl. File 2) revealed that the top enrichment category for cells exposed to osmotic stress and UV-B was iron ion transmembrane transport. However, the direction of change was different for the two stresses; genes associated with iron ion transmembrane transport were upregulated after osmotic stress and downregulated after UV-B stress. One process induced by hyperosmotic stress is vacuolar transport, a response similar to that of plants (Zwiewka et al., 2015). Notably, UV-B stress was associated with the upregulation of RNA-pol I, the RNA polymerase responsible for the transcription of ribosomal genes, and double-strand break repair, while it was associated with the downregulation of key photosynthetic processes.

**Figure 6:**
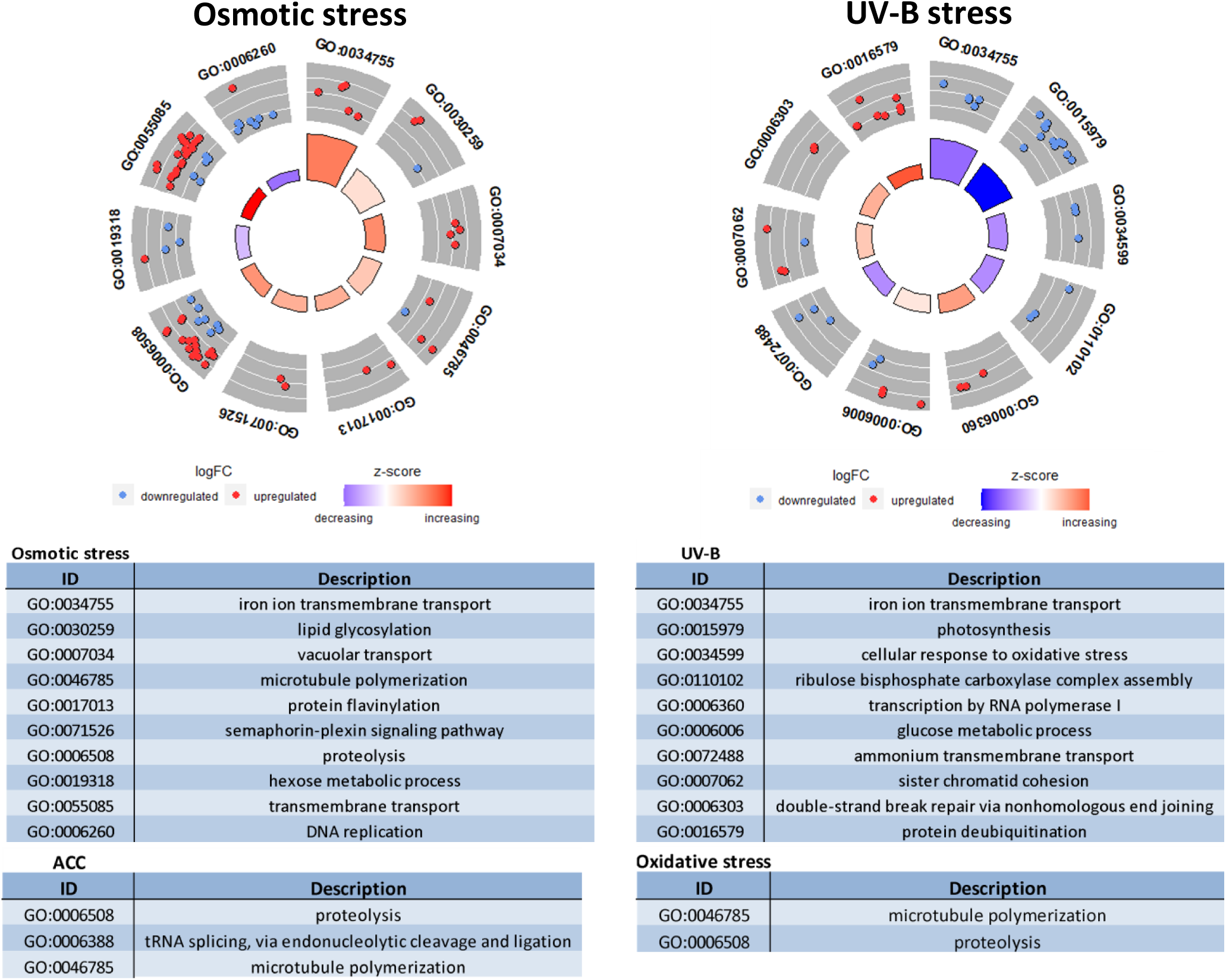
Functional characterization of differentially expressed genes following abiotic stress and ACC treatment. (A) Gene ontology circle plots (GO plots) showing the top significantly enriched GO terms associated with Biological Processes (BP) for differentially expressed genes following abiotic stress. (B) Table displaying the significantly enriched GO terms after ACC treatment.

ACC treatment was associated with a small number of transcriptional changes. Nevertheless, the differentially expressed genes did coalesce around three functional categories: proteolysis, t-RNA splicing, and microtubule polymerization. Similarly, treatment with copper had two enrichment categories: proteolysis and microtubule polymerization.

## DISCUSSION

In this study, we showed that *C. paradoxa,* a primary plastid-bearing alga within the glaucophyte group, produces the hormone ethylene when provided with the plant-like precursor molecule ACC and natively in response to abiotic stress. Additionally, *C. paradoxa* responds to the hormone ABA by producing ROS and with evidence of crosstalk between ABA and ethylene signaling. This is the first report of hormone usage in a glaucophyte alga.

Prior research indicates an early origin of ethylene detection in Archaeplastida, with ethylene receptors likely acquired from a cyanobacterial symbiont, the precursor of the plastid (Gamble et al., 1999). However, the plant-like pathway for ethylene synthesis, using the precursor molecule ACC, appears to have a more recent evolutionary origin (Li et al., 2018; X. Li et al., 2022). This theory is supported by an absence of ACC synthase activity in basal plants and green algae (Xu et al., 2021), the absence of canonical ACC oxidase genes in non-seed-bearing plants (non-spermatophytes) (Li et al., 2023), and an apparent inability of non-spermatophytes to convert ACC to ethylene (Li et al., 2018; X. Li et al., 2022).

Evidence challenging the hypothesis that ACC-based ethylene synthesis originated relatively recently in plant evolution (i.e., in spermatophytes) comes from observations of ACC conversion to ethylene in green and red algae. The charophyte alga *Spirogyra pratensis* converts ACC to ethylene and is reported to have ethylene synthesis and response systems homologous to the land plant version (Ju et al., 2015). Likewise, the green alga *Haematococcus pluvialis* makes ethylene upon ACC exposure (Maillard et al., 1993), and the red alga *Pterocladiella capillacea* produces ethylene natively and reportedly converts ACC into ethylene with an ACO-like activity (García-Jiménez and Robaina, 2012). Our evidence brings this activity to the glaucophytes, further supporting an ancient capacity of the archaeplastid ancestor to produce ethylene when supplied with ACC. In addition to evidence from archaeplastids, amoebozoan slime molds use ethylene to control aspects of the reproductive cycle (Amagai and Maeda, 1992), and inhibitor studies suggest a plant-like pathway of ACC synthesis and conversion to ethylene with an ACO-like enzyme (Amagai, 1987; Amagai et al., 2007).

There is a gap in our knowledge, here, however. While it seems clear that diverse archaeplastids and at least some other protists have the ability to convert ACC to ethylene (although based on available evidence we cannot fully rule out the possibility that ACC induces ethylene formation through an ACC-independent mechanism), native synthesis of ACC itself has not been shown in lineages outside of land plants. Importantly, biochemical tests have demonstrated that ACC-synthase (ACS)-like genes in mosses and green algae, which have sequence and structural homology to plant ACSs, do not have the enzymatic activity that converts S-adenosyl-methionine (SAM) to ACC (X. Li et al., 2022; Sun et al., 2016; Xu et al., 2021). Moreover they lack a second activity common in plant ACS proteins, a carbon-sulfur lyase activity, suggestive of the possibility that plant ACS proteins may have an independent evolutionary origin compared to ACS-like proteins identified by distant homology. All are members of the PLP-dependent enzyme superfamily, with a common domain that may drive homology measures and make bioinformatic-identification of the ancestral function of plant ACS proteins challenging. Despite evidence suggesting that ACS-like proteins in mosses and ferns lack ACS activity, the molecule ACC has been detected in both (Katayose et al., 2021; Li et al., 2018; Xu et al., 2021), suggestive of alternative mechanisms for ACC synthesis in basal plants that may extend to other lineages outside of plants like green algae, red algae, glaucophytes, and amoebozoans: pending measurement of ACC in those organisms and deciphering of mechanisms of synthesis.

ACC can act as a signaling molecule in its own right, independent of its conversion to ethylene (D. Li et al., 2022; Li et al., 2018), causing effects like growth inhibition, even in ethylene insensitive mutant (D. Li et al., 2022; Li et al., 2018). Here we have shown that for the glaucophyte *C. paradoxa* ACC either triggers dose-dependent ethylene production or is directly converted to ethylene in a dose-dependent manner and that ACC treatment causes ROS accumulation and reversible growth inhibition. We also show that treatment with ethephon, a molecule that produces ethylene spontaneously in water independent of ACC, mirrors the effect of ACC treatment, suggesting that even if ACC is a direct signaling molecule, ethylene is at least mirroring its effects in glaucophytes.

ACC conversion to ethylene occurs through the action of the enzyme ACC oxidase (ACO). As noted, organisms from lineages outside of plants, including glaucophytes as reported here, produce ethylene when supplied with exogenous ACC, with one explanation being conversion of ACC to ethylene through an ACO-like protein. The ACO proteins in plants are 2-oxoglutarate and Fe (II)-dependent dioxygenase domain (2OGD)-containing proteins (Kawai et al., 2014) that fall into three phylogenetic clusters, type 1-3 ACOs (Houben and Van de Poel, 2019). Flowering plant ACO proteins are distinct from ACO-like genes in other lineages, including their closest homologs in bryophytes, green algae, and amoebozoans, that may be considered type 4 ACOs given confirmation of their function (Hausinger et al., 2023; Houben and Van de Poel, 2019; Li et al., 2023). Plant ACOs have sequence and structural homology to the bacterial ethylene forming enzyme (EFE) that uses 2-oxoglutarate as a substrate to create ethylene, with the putative type-4 ACOs having a closer affinity to EFE than the seed plant ACOs (type 1-3) (Hausinger et al., 2023). Given the conservation of the active site between canonical ACOs and EFE, one intriguing possibility is that EFE and possibly type-4 ACOs have dual substrate specificity and can make ethylene from both ACC and 2-oxoglutarate, which could explain the presence of ACC oxidase activity in the absence of canonical ACO and ACS genes in lineages outside of flowering plants. To our knowledge this possibility has not been tested biochemically.

Regardless of the mechanism of ACC production, detection, and conversion to ethylene, downstream of ACC we see responses in *C. paradoxa* that mirror some plant responses to stress and ACC/ethylene signaling, such as ROS production and entrance into a quiescence-like state where cell division is inhibited in a reversible fashion and growth can resume once the signal is removed. Halting division is a common stress response in plants and other organisms (Hill et al., 2014; Qi and Zhang, 2020; Sun and Gresham, 2021), presumably as a means to conserve resources and/or avoid damage when organisms are under conditions they are not adapted to. Compared to the broad stress response of *C. paradoxa* to UV and osmotic stress, the response to ACC at the level of gene regulation is minimal, with only 28 upregulated and 9 downregulated genes (Figure 7). That small number of differentially expressed genes, however, has a large apparent effect on the organism. Key genes and protein domains whose expression changes in *C. paradoxa* cells exposed to ACC include an upregulated homolog of the atypical protein kinase RIO1. In the yeast *Saccharomyces cerevisiae*, the protein RIO1 is required for cell cycle progression (Angermayr et al., 2002), and the protein is known to interact with the ribosome, influencing translation (LaRonde-LeBlanc and Wlodawer, 2005). Moreover, a *RIO1* homolog was identified as an upregulated gene under elevated pH, a condition associated with decreased growth, in the diatom *Fragilaria crotonensis* (Zepernick et al., 2022). It is not clear how the upregulation of a protein kinase needed for cell cycle progression influences the observed growth arrest. However, given the specificity of the response to ACC, RIO1 likely plays a significant role in the observed effect. Another upregulated gene encodes an AN1 zinc-finger domain-containing protein known to be involved in stress responses in plants and animals (Vij and Tyagi, 2008). Additionally, a gene encoding Multiprotein Bridging Factor 1 (MBF1) protein was upregulated following ACC exposure. *MBF1* is induced during the plant response to stress (Jaimes-Miranda and Chávez Montes, 2020) and is a bridging factor between transcription factor binding and transcription start sites. At last, four genes encoding subtilisin-like serine proteases (subtilases) were found upregulated. Subtilases have roles in plant biotic and abiotic stress responses and development (Berger and Altmann, 2000; Figueiredo et al., 2017; Schaller et al., 2012). In *C. paradoxa*, the upregulated subtilases may act as general proteases or through specific mechanisms to regulate the cell cycle.

**Figure 7:**
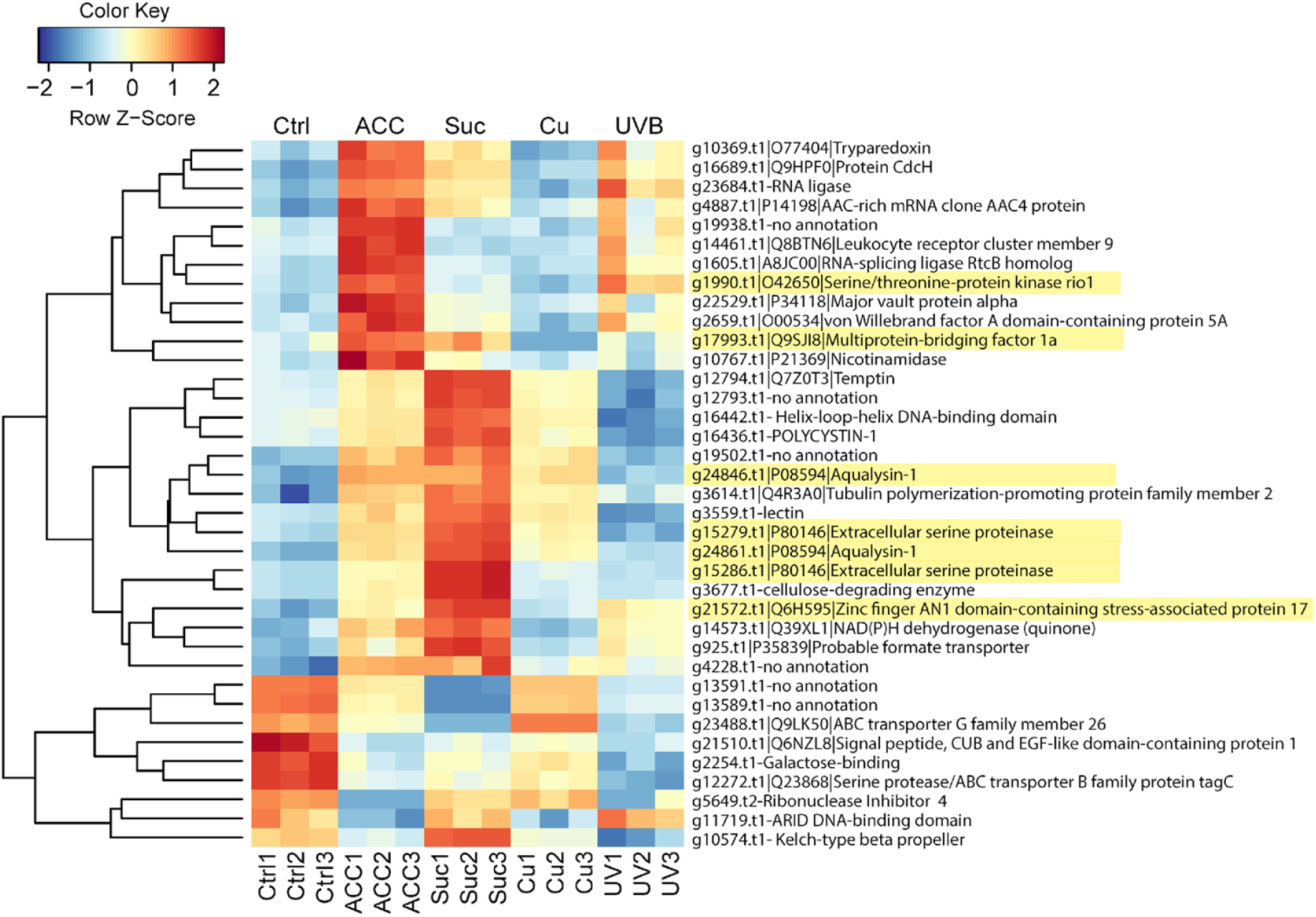
Heatmap of relative expression levels of differentially expressed transcripts associated with ACC treatment across all treatments of *C. paradoxa*. The heatmap shows how genes associated with ACC/ethylene signaling are expressed in control cells and after other treatments. Some expression patterns are shared, while others differ/are unchanged compared to the control in other treatments. Genes discussed in the text are highlighted in yellow (indicated aqualysins and serine proteases are subtilases).

Although ACC exposure has a clear effect on cell division, there is no apparent change in the expression of cyclins at the examined time point (Suppl. Figure 3). This observation contrasts with UV and osmotic stress, which both affect the expression of cyclins, sometimes in opposite directions, with likely effects on cell cycle progression, leading to cells being at different stages of the cell cycle while they recover from the stress (Suppl. Figure 3). For example, Cyclin A2 is downregulated in sucrose-exposed cells at 24 hours but unchanged in UV-exposed cells with respect to the untreated control, suggesting that cells after hyperosmotic stress are not in the S/G2 phase of the cell cycle at the tested time point (Suppl. Figure 3B). Cyclin B2, however, is highly upregulated in UVB-exposed cells and relatively downregulated in sucrose-exposed cells (Suppl. Figure 3B), suggesting that the UVB-exposed cells may be undergoing mitosis at 24 hours, a hypothesis that can be tested in future studies. How the quiescent state is maintained in ACC-exposed cells without changes in expression to cyclin genes compared to the control is not known, though it would be interesting to track control and ACC-exposed cells through time to further examine the role of the cell cycle with respect to the quiescent state induced by ACC.

In addition to the effects of ACC and ethylene, we observed an effect of ABA on ROS production and an additive effect when cells were exposed to both ACC and ABA. However, ABA caused a decrease in the conversion of ACC to ethylene, which could suggest either a decoupling of ACC, ethylene, and ROS production or that the ACC/ethylene concentration was still high enough to have an additive effect. Additional experiments are needed to determine how hormonal crosstalk operates in glaucophytes.

In conclusion, our study on the glaucophyte *C. paradoxa* provides valuable insights into cell signaling via ACC and ethylene in the glaucophyte lineage that may help contextualize the evolution of ethylene signaling in plants (Figure 8). Our results suggest ancient functions of ACC and ethylene signaling that may be shared broadly in the tree of life while in plants ethylene signaling was reinvented when they evolved specific ACS and ACO enzymes that are distinct from their ancient counterparts. The study adds mechanistic insight to the observation that abiotic stress responses are conserved between plants and glaucophytes (Ferrari and Mutwil, 2020) by showing that stress responses may be enacted through hormone signaling pathways in both lineages. An intriguing role for gaseous ethylene acting as a hormone to coordinate stress responses in glaucophytes may be in its ephemeral nature, acting to self-limit the stress response. While glaucophytes appear to make ethylene in response to stress, and ethylene causes growth inhibition, once the stress is removed, gaseous ethylene will diffuse away from the cell and release growth inhibition–as was seen in our experiment where the stress-signal (ACC) was removed and growth resumed (Suppl. Figure 2).

**Figure 8:**
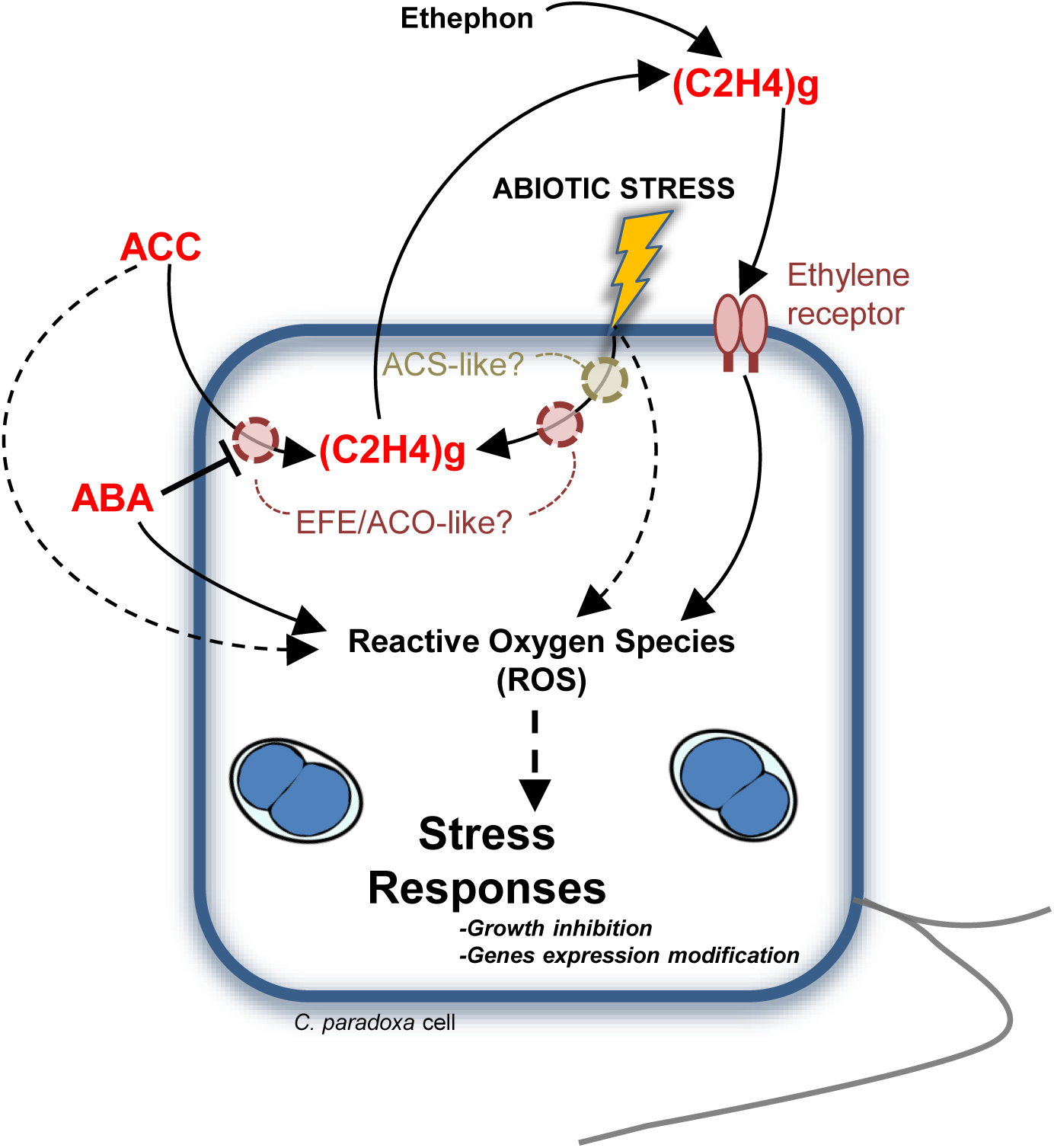
Preliminary model of ethylene signaling in *C. paradoxa*. Schematic representation of the main results of this study. Molecules known as phytohormones are represented in red. Plain black links refer to direct experimental results from this work. Dashed black links represent possible connections in the cell signaling pathway of *C. paradoxa*.

Future studies should aim to identify the specific genes involved in ethylene synthesis and signaling in glaucophytes and explore the potential presence and function of other hormone pathways in other eukaryotes. By doing so, we can gain a more comprehensive understanding of the role and conservation of hormone signaling in the coordination of stress responses and growth regulation across diverse eukaryotes, further our knowledge of the complex interplay between these signaling pathways that have evolved to adapt to various environmental conditions, and establish more robust connections between genomic information and cellular behavior, ultimately developing our understanding of the intricate relationship between genotype and phenotype.

## Data Availability

A mapped read count table for all transcriptome data and the protein structure (.pdb) file from the best Alphafold2 prediction for the glaucophyte 2OGD peptide are available as supplemental data with this manuscript. All raw RNAseq reads were deposited at the NCBI SRA under BioProject #PRJNA979222.

## AKNOWLEDGEMENT

This work was funded in part by the National Science Foundation OIA-1826734. M.G. and M.Y. were supported by a National Science Foundation Research Experience for Undergraduates program at Bigelow Laboratory (NSF EAR 1460861).

## SUPPORTING INFORMATION

<u>Suppl. Figure 1: Ethephon causes ROS accumulation and growth inhibition of *C. paradoxa* cultures.</u>

(A) ROS levels quantification (H2DCFDA staining) 3 days after treatment with 1 mM ethephon and 5 mM ACC. Results were expressed in % increase to control (Mock DMSO). For treatment, ethephon (Sigma 45473) was solubilized in DMSO (300 mM stock) and used to treat *C. paradoxa* cells. For H2DCFDA experiments, ROS were visualized using excitation at 510 nm and a 550 nm emission filter and quantified using a Filter-max-F5 plate reader (Molecular devices).

(B) Fluorescence micrographs of *C. paradoxa* cells treated with 1 mM ethephon and 5 mM ACC. Pictures were taken 3 days after treatments. Fluorescence channels: Red (Chlorophyll) Ex/Em 650/700 nm; Green (H2DCFDA) Ex/Em 510/550 nm. Scale bar represents 350 µm.

(C) Photography of *C. paradoxa* cultures taken 5 days after 1 mM ethephon and 5 mM ACC treatments.

<u>Suppl. Figure 2: ACC-dependent growth inhibition is relieved after washing off ACC from the</u> <u>media.</u>

*C. paradoxa* cell concentrations over 12 days after 5 mM ACC or mock treatments. For each time point, cell concentrations were monitored on day 0, day 5, day 9 and day 12 using a hemocytometer. 20 mL cultures of *C. paradoxa* were treated with 5 mM ACC or mock (water) and incubated in a culture chamber for 5 days. Left panel: At day 5, cells were spin and washed two times in AF6 media before being replaced in 20mL vial and treated with ACC or mock, creating three distinct conditions: ACC + spin/wash in ACC (circle); ACC + spin/wash in mock (triangle); mock + spin/wash in mock (diamond). Right panel: At day 5, cells were not washed and growth was monitored at day 9 and 12. 5 mM ACC treatment (square), mock treatment (triangle). Pictures (bottom): Representatives cultures used for this experiment at day 12 (final point).

<u>Suppl. Figure 3: Heatmap of relative expression levels of differentially expressed transcripts</u> <u>associated with cyclins across all treatments of *C. paradoxa*.</u>

The heatmap shows how genes associated with cyclin are expressed in control cells and after other treatments.

