## Supplemental Figures 1-3 for "Functional stress responses in Glaucophyta: evidence of ethylene and abscisic acid functions in *Cyanophora paradoxa*"

Suppl.Fig1

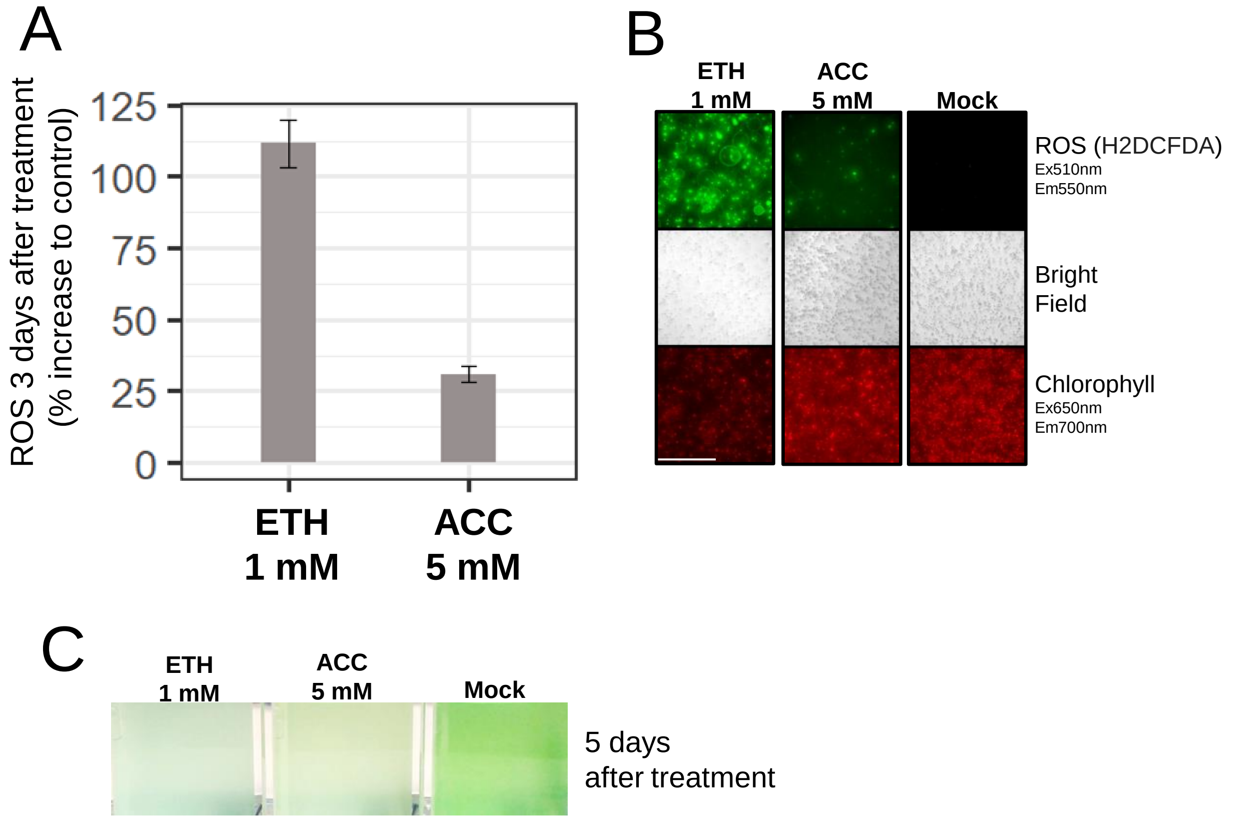
A
ROS 3 days after treatment
(% increase to control)
125
100
75
50
25
0
ETH
1 mM
ACC
5 mM
B
ETH
1 mM
ACC
5 mM
Mock
ROS (H2DCFDA)
Ex510nm
Em550nm
Bright
Field
Chlorophyll
Ex650nm
Em700nm
C
ETH
1 mM
ACC
5 mM
Mock
5 days
after treatment

### Suppl.Fig2

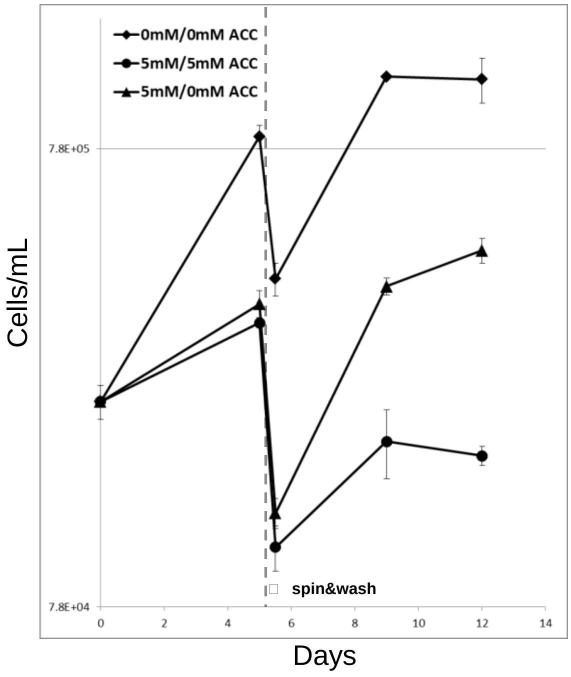


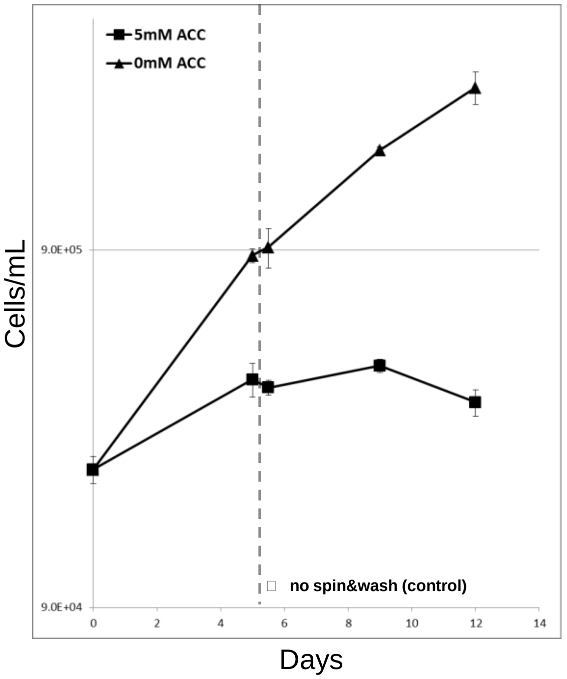


**Day 12 pictures**

**ACC 5mM + spin&wash in AF6** *5mM/0mM ACC*

**ACC 5mM + spin&wash in ACC 5mM** *5mM/5mM ACC*

**Mock + spin&wash in AF6** *0mM/0mM ACC*

**Mock** **ACC 5mM**

**no spin&wash**

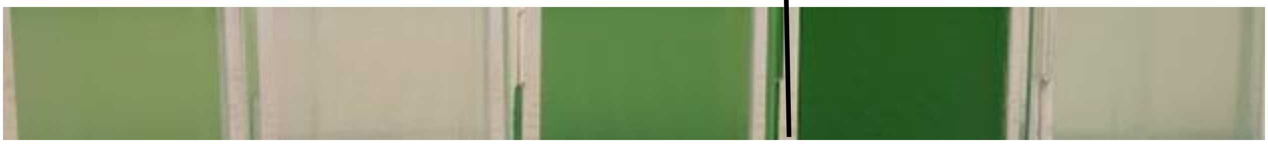

**Suppl.Fig3**

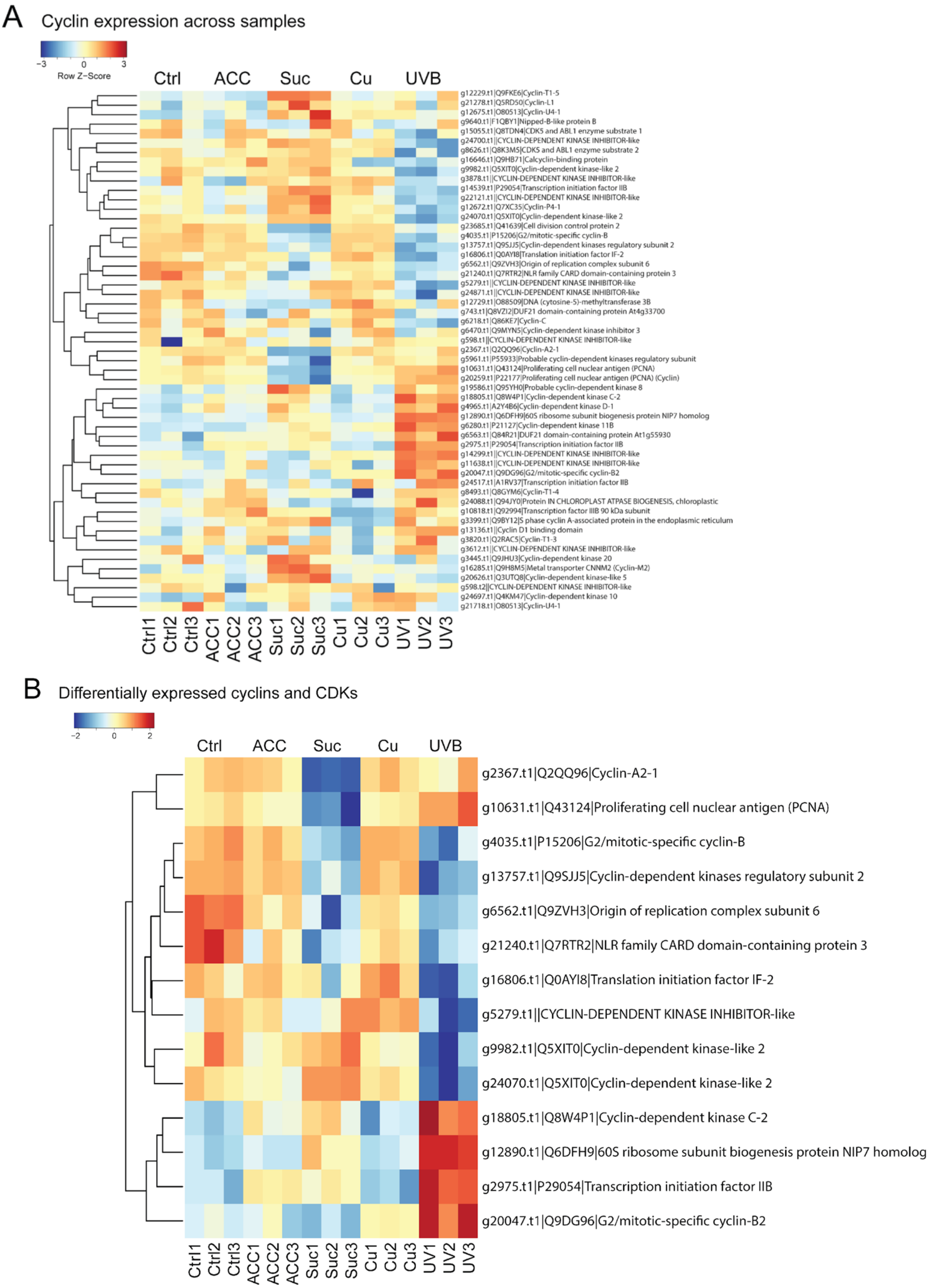
A
Cyclin expression across samples
−3
0
3
Row Z−Score
Ctrl
ACC
Suc
Cu
UVB
g12229.t1|Q9FKE6|Cyclin-T1-5
g21278.t1|Q5RD50|Cyclin-L1
g12675.t1|O80513|Cyclin-U4-1
g9640.t1|F1QBY1|Nipped-B-like protein B
g15055.t1|Q8TDN4|CDK5 and ABL1 enzyme substrate 1
g24700.t1||CYCLIN-DEPENDENT KINASE INHIBITOR-like
g8626.t1|Q8K3M5|CDK5 and ABL1 enzyme substrate 2
g16646.t1|Q9HB71|Calcyclin-binding protein
g9982.t1|Q5XIT0|Cyclin-dependent kinase-like 2
g3878.t1||CYCLIN-DEPENDENT KINASE INHIBITOR-like
g14539.t1|P29054|Transcription initiation factor IIB
g22121.t1||CYCLIN-DEPENDENT KINASE INHIBITOR-like
g12672.t1|Q7XC35|Cyclin-P4-1
g24070.t1|Q5XIT0|Cyclin-dependent kinase-like 2
g23685.t1|Q41639|Cell division control protein 2
g4035.t1|P15206|G2/mitotic-specific cyclin-B
g13757.t1|Q9SJJ5|Cyclin-dependent kinases regulatory subunit 2
g16806.t1|Q0AYI8|Translation initiation factor IF-2
g6562.t1|Q9ZVH3|Origin of replication complex subunit 6
g21240.t1|Q7RTR2|NLR family CARD domain-containing protein 3
g5279.t1||CYCLIN-DEPENDENT KINASE INHIBITOR-like
g24871.t1||CYCLIN-DEPENDENT KINASE INHIBITOR-like
g12729.t1|O88509|DNA (cytosine-5)-methyltransferase 3B
g743.t1|Q8VZI2|DUF21 domain-containing protein At4g33700
g6218.t1|Q86KE7|Cyclin-C
g6470.t1|Q9MYN5|Cyclin-dependent kinase inhibitor 3
g598.t1||CYCLIN-DEPENDENT KINASE INHIBITOR-like
g2367.t1|Q2QQ96|Cyclin-A2-1
g5961.t1|P55933|Probable cyclin-dependent kinases regulatory subunit
g10631.t1|Q43124|Proliferating cell nuclear antigen (PCNA)
g20259.t1|P22177|Proliferating cell nuclear antigen (PCNA) (Cyclin)
g19586.t1|Q95YH0|Probable cyclin-dependent kinase 8
g18805.t1|Q8W4P1|Cyclin-dependent kinase C-2
g4965.t1|A2Y4B6|Cyclin-dependent kinase D-1
g12890.t1|Q6DFH9|60S ribosome subunit biogenesis protein NIP7 homolog
g6280.t1|P21127|Cyclin-dependent kinase 11B
g6563.t1|Q84R21|DUF21 domain-containing protein At1g55930
g2975.t1|P29054|Transcription initiation factor IIB
g14299.t1||CYCLIN-DEPENDENT KINASE INHIBITOR-like
g11638.t1||CYCLIN-DEPENDENT KINASE INHIBITOR-like
g20047.t1|Q9DG96|G2/mitotic-specific cyclin-B2
g24517.t1|A1RV37|Transcription initiation factor IIB
g8493.t1|Q8GYM6|Cyclin-T1-4
g24088.t1|Q94JY0|Protein IN CHLOROPLAST ATPASE BIOGENESIS, chloroplastic
g10818.t1|Q92994|Transcription factor IIIB 90 kDa subunit
g3399.t1|Q9BY12|S phase cyclin A-associated protein in the endoplasmic reticulum
g13136.t1||Cyclin D1 binding domain
g3820.t1|Q2RAC5|Cyclin-T1-3
g3612.t1||CYCLIN-DEPENDENT KINASE INHIBITOR-like
g3445.t1|Q9JHU3|Cyclin-dependent kinase 20
g16285.t1|Q9H8M5|Metal transporter CNNM2 (Cyclin-M2)
g20626.t1|Q3UTQ8|Cyclin-dependent kinase-like 5
g598.t2||CYCLIN-DEPENDENT KINASE INHIBITOR-like
g24697.t1|Q4KM47|Cyclin-dependent kinase 10
g21718.t1|O80513|Cyclin-U4-1
Ctrl1
Ctrl2
Ctrl3
ACC1
ACC2
ACC3
Suc1
Suc2
Suc3
Cu1
Cu2
Cu3
UV1
UV2
UV3
B
Differentially expressed cyclins and CDKs
−2
0
2
Ctrl
ACC
Suc
Cu
UVB
g2367.t1|Q2QQ96|Cyclin-A2-1
g10631.t1|Q43124|Proliferating cell nuclear antigen (PCNA)
g4035.t1|P15206|G2/mitotic-specific cyclin-B
g13757.t1|Q9SJJ5|Cyclin-dependent kinases regulatory subunit 2
g6562.t1|Q9ZVH3|Origin of replication complex subunit 6
g21240.t1|Q7RTR2|NLR family CARD domain-containing protein 3
g16806.t1|Q0AYI8|Translation initiation factor IF-2
g5279.t1||CYCLIN-DEPENDENT KINASE INHIBITOR-like
g9982.t1|Q5XIT0|Cyclin-dependent kinase-like 2
g24070.t1|Q5XIT0|Cyclin-dependent kinase-like 2
g18805.t1|Q8W4P1|Cyclin-dependent kinase C-2
g12890.t1|Q6DFH9|60S ribosome subunit biogenesis protein NIP7 homolog
g2975.t1|P29054|Transcription initiation factor IIB
g20047.t1|Q9DG96|G2/mitotic-specific cyclin-B2
Ctrl1
Ctrl2
Ctrl3
ACC1
ACC2
ACC3
Suc1
Suc2
Suc3
Cu1
Cu2
Cu3
UV1
UV2
UV3
